# Proof of concept: Targeted protein degradation of the stress granules component G3BP1 as an antiviral strategy against norovirus infection

**DOI:** 10.1101/2025.07.21.666062

**Authors:** Liliana Echavarria-Consuegra, Ian Goodfellow

**Affiliations:** Division of Virology, Department of Pathology, University of Cambridge, Cambridge, United Kingdom

**Keywords:** G3BP1, PROTAC, targeted-protein-degradation, Norovirus, Antiviral

## Abstract

Human norovirus (HNoV) is a major cause of gastroenteritis worldwide, for which no antiviral therapies exist to date. Previously, our lab has demonstrated that both HNoV and murine norovirus (MNV1) are highly dependant on the expression of the Ras-GTPase-activating protein-binding protein 1 (G3BP1), a cellular protein mostly involved in the assembly of stress granules. We therefore hypothesize that targeting G3BP1 could be a promising antiviral strategy against noroviruses. Here, we designed a proof-of concept study to test targeted protein degradation as a mechanism to induce the specific proteolysis of G3BP1 via the proteasome. To do so, we generated a cellular platform for the over-expression of G3BP1 fused to the bacterial protein Halotag (HaloG3BP1). First, we showed that MNV1 replication is restored in G3BP1-knockout (ΔG3BP1) cells complemented with HaloG3BP1. We then used a proteolysis-targeting-chimera directed towards the Halotag (HaloPROTAC), to induce the specific degradation of HaloG3BP1. We further demonstrate that proteolysis of G3BP1 reduces MNV1 replication, leading to a lower infectious virus yield and preventing virus-induced cell death. We also confirmed that the mechanism of HaloPROTAC3 is mediated via the recruitment of Cullin2-VHL E3-ubiquitin ligase. Our findings add to the body of evidence supporting that targeting of the cellular protein G3BP1 can be used as an antiviral approach, and validates the use of PROTACs for the efficient and specific degradation of cellular factors as a feasible methodology to combat viral diseases.

## Introduction

Human norovirus (HNoV) is a significant cause of global morbidity in the human population, causing ∼685 million cases of acute gastroenteritis yearly (1, 2). While HNoV typically leads to self-limiting illness in the immunocompetent, infection can result in chronic disease and even death in high-risk groups, such as young children and the elderly (2–4). Outbreaks caused by HNoV are commonly associated with semi-enclosed settings such as hospitals wards and nursing homes (5). The high risk of transmission in these settings is mainly attributed to the low infectious dose of HNoV and the extended shedding period, highlighting the potential need for prophylactic antivirals to control virus spread in vulnerable groups (6).

HNoV is a non-enveloped virus with a single-stranded RNA genome of positive polarity that is classified within the genus *Norovirus* of the *Caliciviridae* family (7, 8). Noroviruses are characterised by an icosahedral capsid containing the RNA genome, which ranges in size from 7.4 to 7.7 Kb and is organised in 3–4 open reading frames (ORFs) (9). A large polyprotein containing 6 non-structural proteins is encoded within ORF1, whereas ORF2 and 3 encode the major and minor capsid proteins, respectively (9). Murine norovirus (MNV1) genome contains a fourth ORF overlapping ORF2, which produces an accessory protein known as virulence factor 1 (VF1) (10). The norovirus genome is covalently linked at the 5’ end to the viral protein VPg (11), which functions as a proteinaceous 5’ cap substitute, recruiting translation initiation factors to drive viral translation in a VPg-dependent manner (12–14).

Despite its clinical significance, antiviral treatments or vaccines specifically designed to target HNoV are currently unavailable. One of the biggest challenges hindering drug discovery is the lack of a cost-effective and robust cell culture system that supports the full life cycle of HNoV (15, 16). Recent breakthroughs allowing HNoV propagation *in vitro*, such as cultured B cells and human intestinal enteroids, while valuable, pose significant technical challenges associated with cost and reproducibility in order to be readily suitable for large scale antiviral screens (17–20). Other alternatives such as Norwalk virus replicon-bearing cell lines or surrogate viruses from within the *Caliciviridae* family have been exploited to study antiviral candidates (21). Among the latter, MNV1 has been widely used, since it belongs to the same genus as HNoV and shares tropism for immune and intestinal epithelial cells (22).

Preclinical antiviral studies for noroviruses have mostly focused on conventional therapies that directly target specific steps of the virus life cycle or the viral proteins (23). One of the major disadvantages of direct-acting antivirals, however, is the emergence of resistance mutations, which reduces their potential use. Research on host-targeted therapeutics has emerged as an alternative, as these molecules often have a higher barrier to resistance and could potentially target proteins that are relevant for multiple stages of the virus life cycle (23–26). Thus, several host factors with essential roles in the norovirus life cycle represent promising candidates for host-targeted antiviral intervention. We and others have described, for example, that the host protein Ras GTPase-activating protein-binding protein 1 (G3BP1) is essential for the leading rounds of norovirus protein translation and virus genome replication (27–29). G3BP1 is a multifunctional protein, better known for its role as a molecular switch that drives stress granules assembly during translational shutoff, due to the cytoplasmic accumulation of released mRNA molecules from polysomes (30, 31). Stress granules are dynamic condensates, where mRNA is temporarily stored until stress is resolved and the cellular environment returns to homeostatic conditions (32). While norovirus infection is accompanied by eIF2α phosphorylation and consequently host translational arrest, G3BP1 is redistributed to virus replication complexes, favouring virus protein translation and replication through a mechanism not yet fully understood (27, 28). In line with this, genetic ablation of G3BP1 significantly disrupts the replication of both MNV1 and HNoV (27, 29). This evidence suggests that targeting G3BP1 may hold future therapeutic potential against noroviruses.

Host-targeted therapeutics can be achieved using targeted protein degradation, which has emerged as a novel strategy to induce chemical knockdown of a protein of interest (POI), in contrast to the traditional regulation of protein activity using small molecule inhibitors (33). Modulation of a POI through degradation can be accomplished using PROteolysis TArgeting Chimeras (PROTACs), heterobifunctional molecules composed of a ligand for the POI and a ligand for an E3 ubiquitin ligase (34). Simultaneous binding to both targets, leads to POI ubiquitination and subsequent degradation by the ubiquitin-proteasome system (34). In antiviral drug discovery, PROTACs have been developed in proof-of-concept studies to successfully target a wide variety of viruses, including hepatitis B virus, hepatitis C virus, Influenza A virus and SARS-CoV-2 (35). More recently, a few PROTACs targeting human factors have been explored as antivirals against coronaviruses, HCV and human immunodeficiency virus (36, 37).

To establish proof of concept that PROTAC-mediated degradation of G3BP1 can serve as an antiviral strategy against noroviruses, we first engineered a cellular model in which Halotag-fused G3BP1 could be selectively targeted for proteasomal degradation using commercially available HaloPROTACs. PROTACs targeting the Halotag (HaloPROTACs) were then used to drive the degradation of Halo-tagged G3BP1, as it has been previously done for other cellular and exogenous targets (38).

Here, we show that HaloPROTAC-mediated degradation of G3BP1 leads to significantly reduced norovirus replication using the MNV1 infection model. We found that G3BP1 is bound to and degraded by HaloPROTAC3 in a dose-dependent manner, leading to a reduction in the target protein levels in the model. Furthermore, we confirmed that G3BP1 chemical ablation, using this specific degrader, led to a clear reduction in the levels of norovirus RNA and infectious yield levels, and that this function was mediated by the ubiquitin-proteasome system. This work adds to the growing body of evidence on the potential value of PROTAC degraders as a therapeutic approach against viruses.

## Results

### Development of a model for stable HaloG3BP1 expression in BV2 ΔG3BP1 cells

Norovirus VPg-dependant protein translation and therefore subsequent virus replication, rely heavily on the expression of the cellular protein G3BP1. Using western blot analysis and TCID_50_, we confirmed that MNV1 fails to produce viral proteins and infectious progeny when G3BP1 is ablated (Figure 1A-B). Due to its role in norovirus life cycle, we reasoned G3BP1 to be a valuable antiviral target for noroviruses. We therefore sought to develop a cellular model in which G3BP1 could be targeted for proteasomal degradation using commercially available HaloPROTACs as, at the time this study was initiated, no validated G3BP1-targeting PROTACs, were available. To that aim, human G3BP1 fused to the C-terminal of Halotag, a modified bacterial dehalogenase, was subcloned into a lentiviral vector under the control of the human phosphoglycerate kinase (hPGK) promoter. The lentiviral vector used was bicistronic and contained a hygromycin resistance gene under the translational control of HCMV IRES. Infectious lentiviruses were used to transduce BV2 cells, either parental wild-type (WT) cells or those lacking G3BP1 (ΔG3BP1), and stable cells were generated by hygromycin B selection. As controls, BV2 WT and ΔG3BP1 cells were transduced with lentiviruses encoding the Halotag alone. High Halotag-expressing cells were enriched by cell sorting and the expression of the Halotag or Halotag-G3BP1 fusion protein (HaloG3BP1) were confirmed by western blot (Figure 1C). As expected, HaloG3BP1 showed much higher expression levels than endogenous G3BP1 due to the strong activity of the hPGK promoter. Of note, we observed that the expression of Halotag alone in BV2 and ΔG3BP1 cells was higher than that of the fused HaloG3BP1 protein.

**Figure 1.**
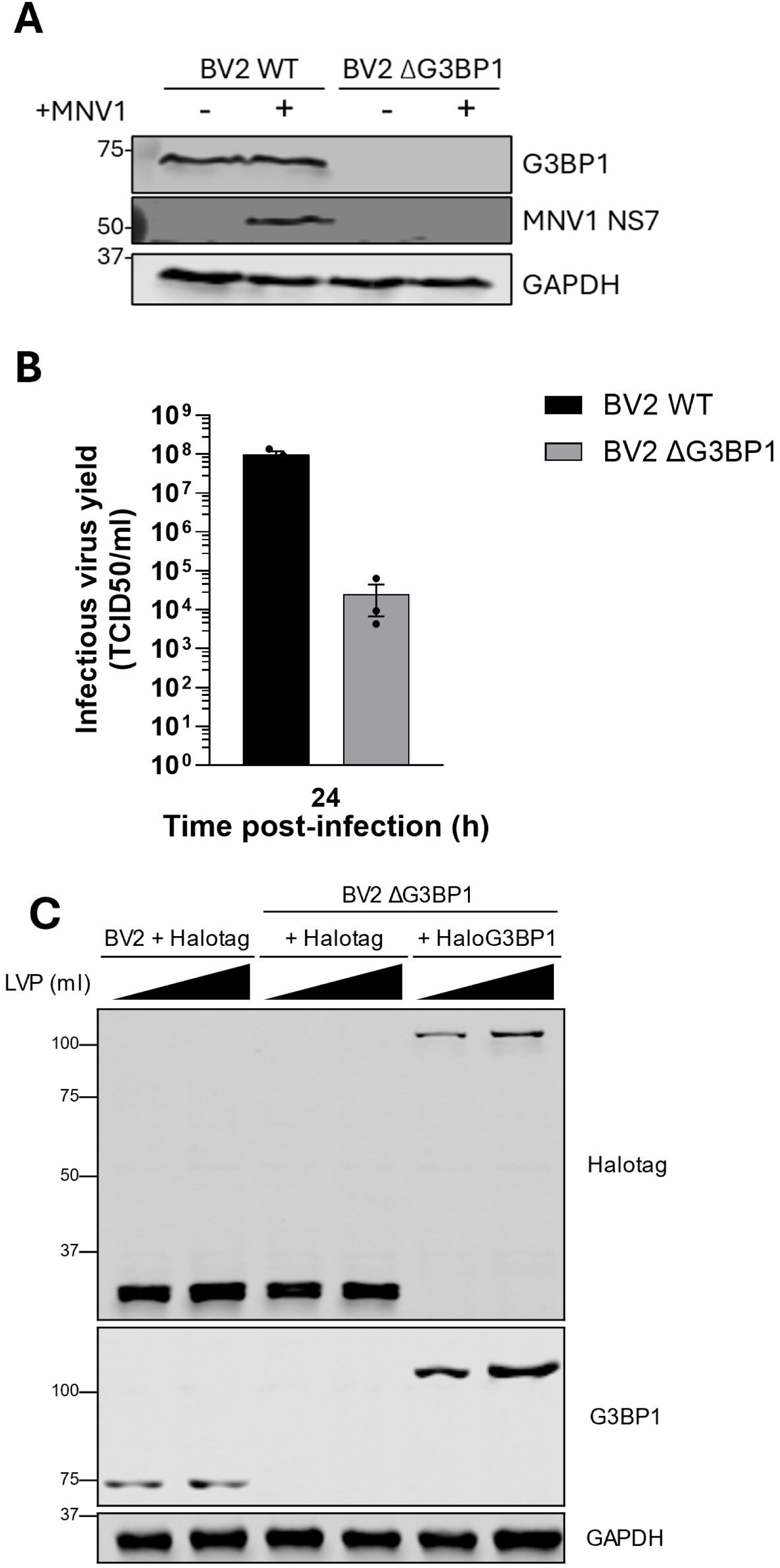
G3BP1 ablation impairs norovirus infection and generation of HaloG3BP1-expressing cells. BV2 WT or ΔG3BP1 cells were infected with an MOI of 10 TCID_50_/cell of MNV1. (A) After 8 h.p.i., protein samples were collected for western blot analysis with antibodies targeting G3BP1, MNV1 NS7 and GAPDH. (B) Total virus infectious yield after 24 h.p.i. was quantified by TCID_50_ in BV2 WT cells. (C) Protein samples from BV2 WT or ΔG3BP1 cells transduced with either 1 or 2 ml of lentivirus-containing supernatant (LVP) were subjected to western blot analysis using antibodies directed against Halotag, G3BP1 or GAPDH. Endogenous G3BP1 protein in BV2 + Halotag cells has an observed molecular weight of 68 kDa, whereas HaloG3BP1 fusion protein has a calculated molecular weight of 101 kDa.

To confirm that norovirus replication was rescued in ΔG3BP1 cells reconstituted with HaloG3BP1, cells were infected with MNV1 and infection levels were evaluated. Robust norovirus replication and virus progeny release in BV2 cells is accompanied by extensive cell death, we therefore used cell death as a proxy of active infection. Cells were infected with MOI 10, 1 and 0.1 TCID_50_/cell and virus-induced cell death was followed in real time using the live-exclusion dye NIR in an Incucyte imaging system (Figure 2A and B). BV2 cells infected with MNV1 showed extensive virus-induced cell death (NIR+ cell count) as early as 12 h.p.i., when infected with MNV1 at MOI 10 TCID_50_/cell (Figure 2B). Infection with MOI 1 and 0.1 TCID_50_/cell led to extensive cell death after ∼15 and ∼18 h.p.i., respectively. Expression of the Halotag had no effect on MNV1-induced cell death, as BV2 + Halotag cells had similar levels of NIR+ counts to that of BV2 WT cells. As expected, virus replication in BV2 lacking G3BP1 was significantly impaired, and the expression of the Halotag alone had no impact on this phenotype. In contrast, MNV1-induced cell death was evident after 21 h.p.i. in BV2 ΔG3BP1 + HaloG3BP1 cells, in agreement with a restoration of the function of G3BP1 in norovirus infection. The observed minor delay in cell death induction in comparison to BV2 WT cells, was likely a result of an effect of the Halotag on G3BP1 function or due to the high expression levels having a negative impact on protein functionality. The ability of MNV1 to infect and induce cell death was also assessed in an end-point cytolysis assay (Figure 2C). In agreement with our previous observations, while low doses of MNV1 (∼1 TCID_50_/well) were sufficient to kill BV2 WT and BV2 + Halotag cells after 5 d.p.i., a much higher infectious dose, i.e. 10^4^ to 10^3^ TCID_50_/well, was required to kill cells where G3BP1 was ablated, irrespective of the expression of Halotag. BV2 ΔG3BP1 cells that were reconstituted with HaloG3BP1 were, in contrast, much more susceptible to low doses of MNV1, and showed virus-induced cell death, requiring only ∼10 TCID_50_/well to cause complete monolayer destruction at 5 d.p.i. (Figure 2C).

**Figure 2.**
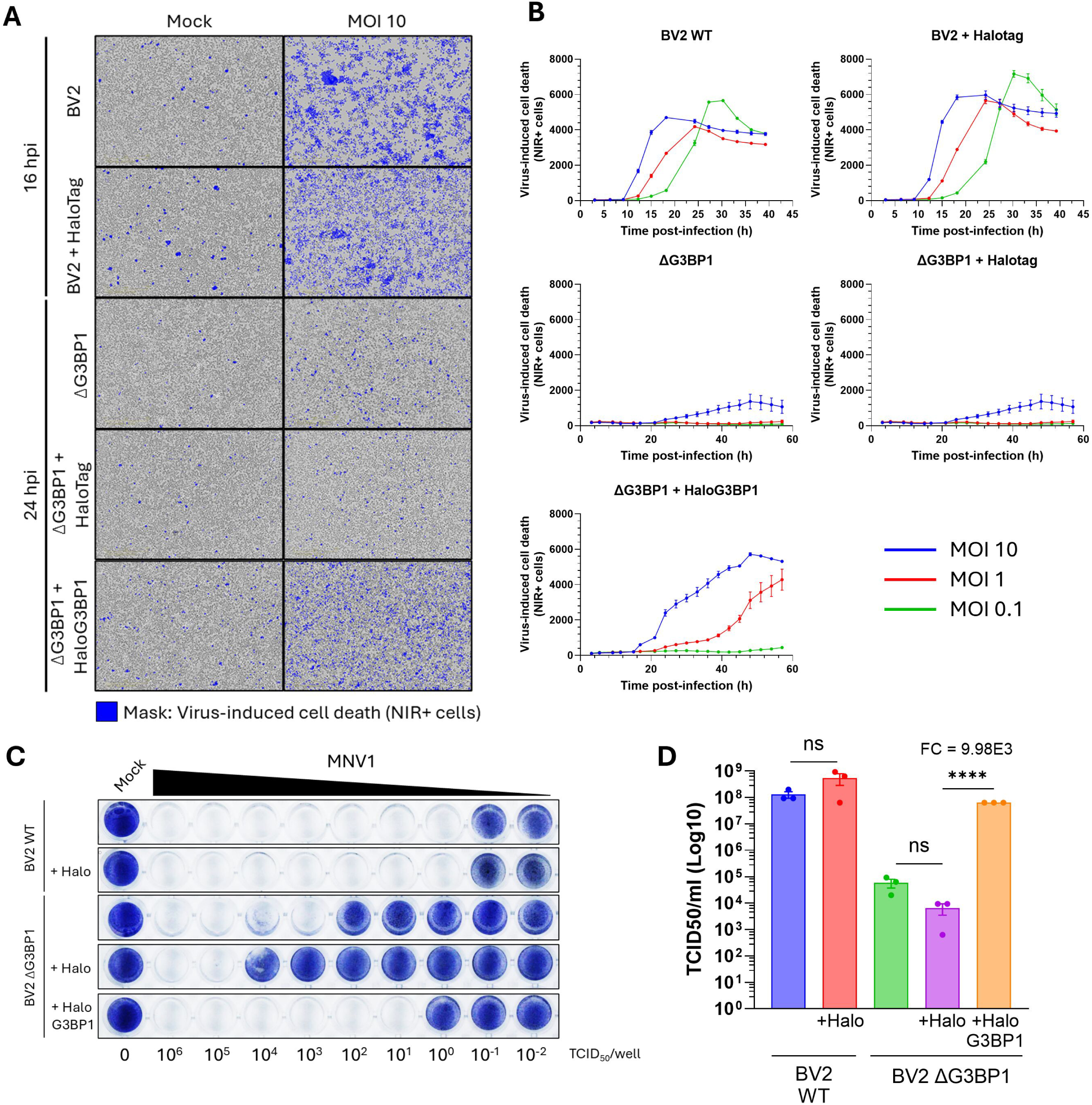
HaloG3BP1 rescues MNV1 infection in BV2 ΔG3BP1 cells. BV2, BV2 + Halotag, ΔG3BP1, ΔG3BP1 + Halotag or ΔG3BP1 + HaloG3BP1 cells were infected with MNV1 at the indicated MOI. (A-B) Infection-induced cell death was detected using the NIR dye measured in the live cell imaging Incucyte system. (A) Representative pictures after 16 or 24 h.p.i., NIR + cells are shown in blue. (B) Quantification of the total number of NIR + cells in all the different cell lines was measured every 3 h and followed for a total of 60 h.p.i. (C) The indicated cell lines were plated in a 96 well plate and subsequently infected using a 10-fold serial dilution of MNV1, starting at 10^6^ TCID_50_/well. Cells were fixed in paraformaldehyde and stained with toluidine blue 5 d.p.i. to evidence cell death. (D) Total virus infectious yield after 24 h.p.i. was quantified by TCID_50_ in BV2 WT cells.

Lastly, to confirm norovirus replication in cells expressing the fusion HaloG3BP1 protein, MNV1 infectious yield was quantified by TCID_50_. Infection of BV2 or BV2 + Halotag cells resulted in an infectious titre of ∼10^8^ TCID_50_/ml after 24 h.p.i. (Figure 2D). Conversely, in the absence of G3BP1 and in agreement with our previous observations, the infectious virus yield was reduced by ∼10,000-fold in comparison to BV2 WT cells. However, restoration of G3BP1 expression in the form of HaloG3BP1 restored replication to near WT levels (Figure 2D).

### HaloPROTAC3 binds HaloG3BP1 fusion protein inducing its degradation

Having established a cellular model suitable for targeting G3BP1 using HaloPROTACs, we selected HaloPROTAC3 in the first instance, as it has proven to effectively degrade Halotag-fusion proteins (38, 39). HaloPROTAC3 is derived from a chloroalkane moiety targeting Halotag and VH285, a ligand for Cullin2-VHL E3 ubiquitin ligase (Figure 3A). The cytotoxicity of HaloPROTAC3 was initially determined over a range of concentrations (Figure 3B), after 48 h post-treatment (h.p.t.) (CC_50_ = 16.42 µM). Concentrations of HaloPOTAC3 below 2 µM resulted in less than ∼30% cell death and were therefore selected for follow-up studies. First, to investigate HaloPROTAC3 binding and degradation of Halotag or HaloG3BP1 via its chloroalkane moiety, a flow cytometry assay using a Halotag fluorescent ligand (Janelia 647) was carried out. The fluorescent ligand used binds Halotag or HaloG3BP1 molecules that are not engaged by HaloPROTAC3, thereby simultaneously providing a readout that combines target engagement by the HaloPROTAC3 and cellular expression levels of the target. Figure 3C shows the gating strategy, depicting the exclusion of dead cells from the analysis by staining with Live/Dead (UV) viability dye. Representative histograms of the Halotag fluorescence in the Janelia 647 channel for the DMSO control or cells treated with decreasing concentrations of HaloPROTAC3 are shown in Figure 3D. The percentage of Halotag positive (Halotag +) cells after 48 h.p.t. in the DMSO control samples was ∼97-98% (Figure 3D-E). Halotag expression in BV2 and ΔG3BP1 cells exhibited a bimodal distribution, indicating the presence of two sub-populations with either dim or bright Halotag expression, likely due to the polyclonal nature of the cells used in this study (Figure 3D). The expression of HaloG3BP1 fusion protein displayed a more discrete distribution, resulting in one unique population of Halotag + cells. In all cases, treatment with HaloPROTAC3 resulted in a dose-dependent decrease in the percentage of Halotag + cells in comparison with cells treated with DMSO, confirming target engagement and likely protein degradation. Cells treated with HaloPROTAC3 concentrations in between 1.2 µM and 600 nM showed a striking reduction in the number of cells expressing Halotag or HaloG3BP1, whereas concentrations below 150 nM failed to cause any detectable changes in the number of cells positive for the target in this assay (Figure 3D-E). The Halotag mean fluorescence intensity (MFI) from these experiments was used to estimate the concentration of HaloPROTAC3 resulting in half of its maximum achievable degradation (DC_50_) in the various cell lines (Figure 3E). We note, however, that the probe and HaloPROTAC3 share binding sites, such that any change in MFI represents a combination of degradation and target engagement. Noting these caveats, we estimated a DC_50_ of 641.9 nM in BV2 + Halotag cells, 852.6 nM in ΔG3BP1 + Halotag cells and 632.1 nM in ΔG3BP1 + HaloG3BP1 cells (Figure 3E).

**Figure 3.**
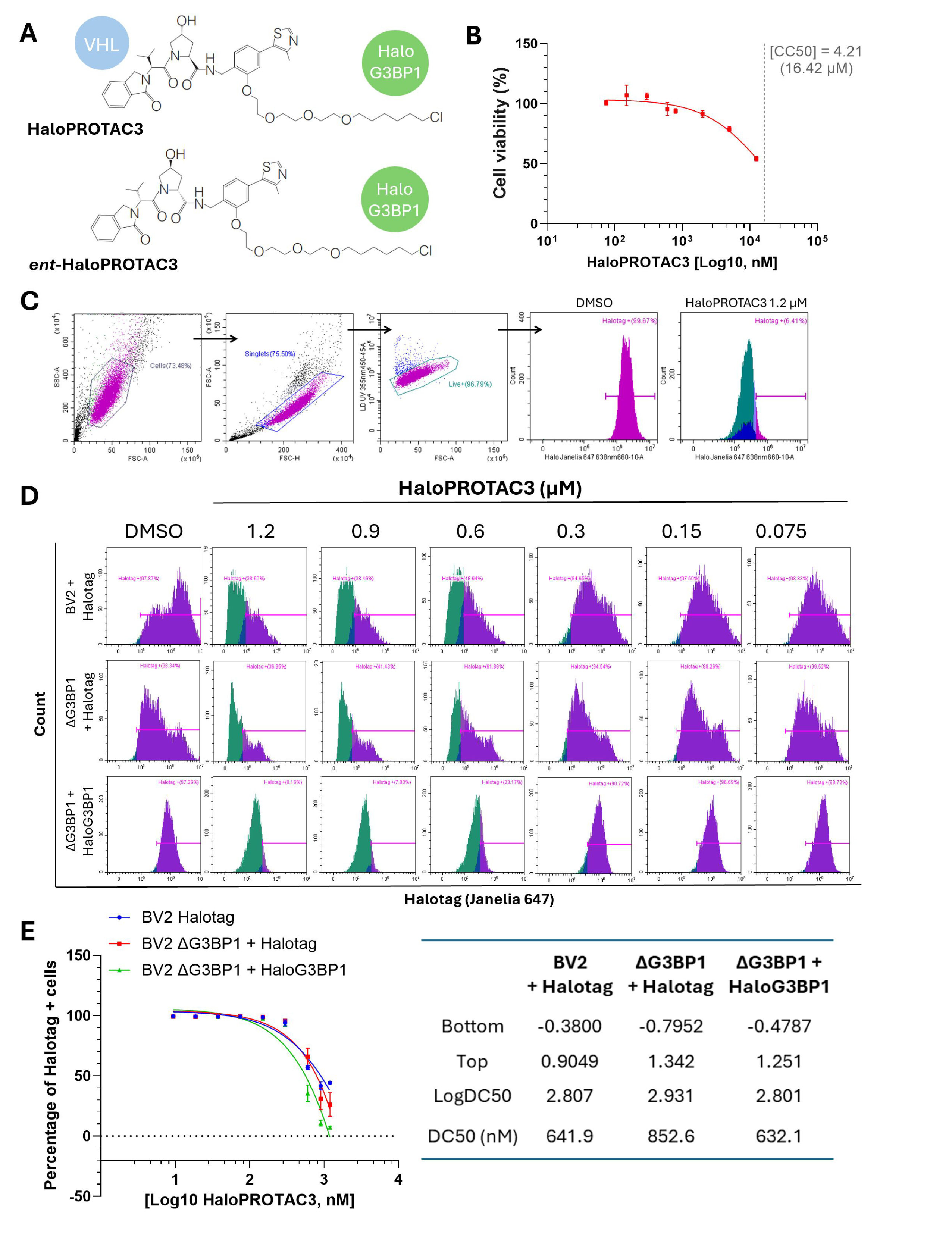
HaloPROTAC3 efficiently binds Halotag and HaloG3BP1 fusion proteins. (A) Chemical structure of HaloPROTAC3 and the enantiomeric form *ent*-HaloPROTAC3, scheme indicates the warhead binding VHL E3 ubiquitin ligase (VH285) and the chloroalkene moiety binding Halotag. (B) BV2 ΔG3BP1 + HaloG3BP1 cells were treated with the indicated concentrations of HaloPROTAC3 for 48 h, and metabolic cell death was measured using Cell titer blue. Cell death was calculated as relative to the DMSO control and CC_50_ is indicated by a dotted line. (C-E) BV2 + Halotag, BV2 ΔG3BP1 + Halotag or BV2 ΔG3BP1 + HaloG3BP1 cells treated with the indicated concentrations of HaloPROTAC3 for 48 h were subjected to flow cytometry analysis after incubation for 2 h with the fluorescent ligand Halotag Janelia 647. (C) Gating strategy of Halotag + cells in the live cell population. (D) Representative histograms and (E) Dose-response curves of the quantified percentage of Halotag + cells. Shown, are the corresponding curve fitting parameters for each cell line.

To confirm that Halotag and HaloG3BP1 engagement by HaloPROTAC3 leads to protein degradation in an orthogonal assay, Halotag and G3BP1 protein levels in cells treated with HaloPROTAC3 were assessed by western blot. In agreement with the flow cytometry data, HaloPROTAC3 showed to effectively bind Halotag in cells after 24 h.p.t. (Figure 4A), as suggested by the apparent change in protein size in comparison to the non-treated cells, indicative of the covalent binding of this degrader to its target. Of note, this shift in mass was not observed in ΔG3BP1 cells expressing HaloG3BP1 due to the much larger mass of the fusion protein and the inability to resolve the different species on the SDS-PAGE gel. Protein degradation was observed from 24 h onwards (Figure 4A-B), with degradation continuing to increase after 48 h.p.t., when protein levels were decreased to ∼39% and ∼22% in BV2 + Halotag and ΔG3BP1 + Halotag cells respectively. In the case of BV2 ΔG3BP1 + HaloG3BP1 cells, protein levels were visibly reduced after 24 h.p.t (∼45% in comparison to the control), which further decreased after 48 h.p.t. to ∼6% of the control.

**Figure 4.**
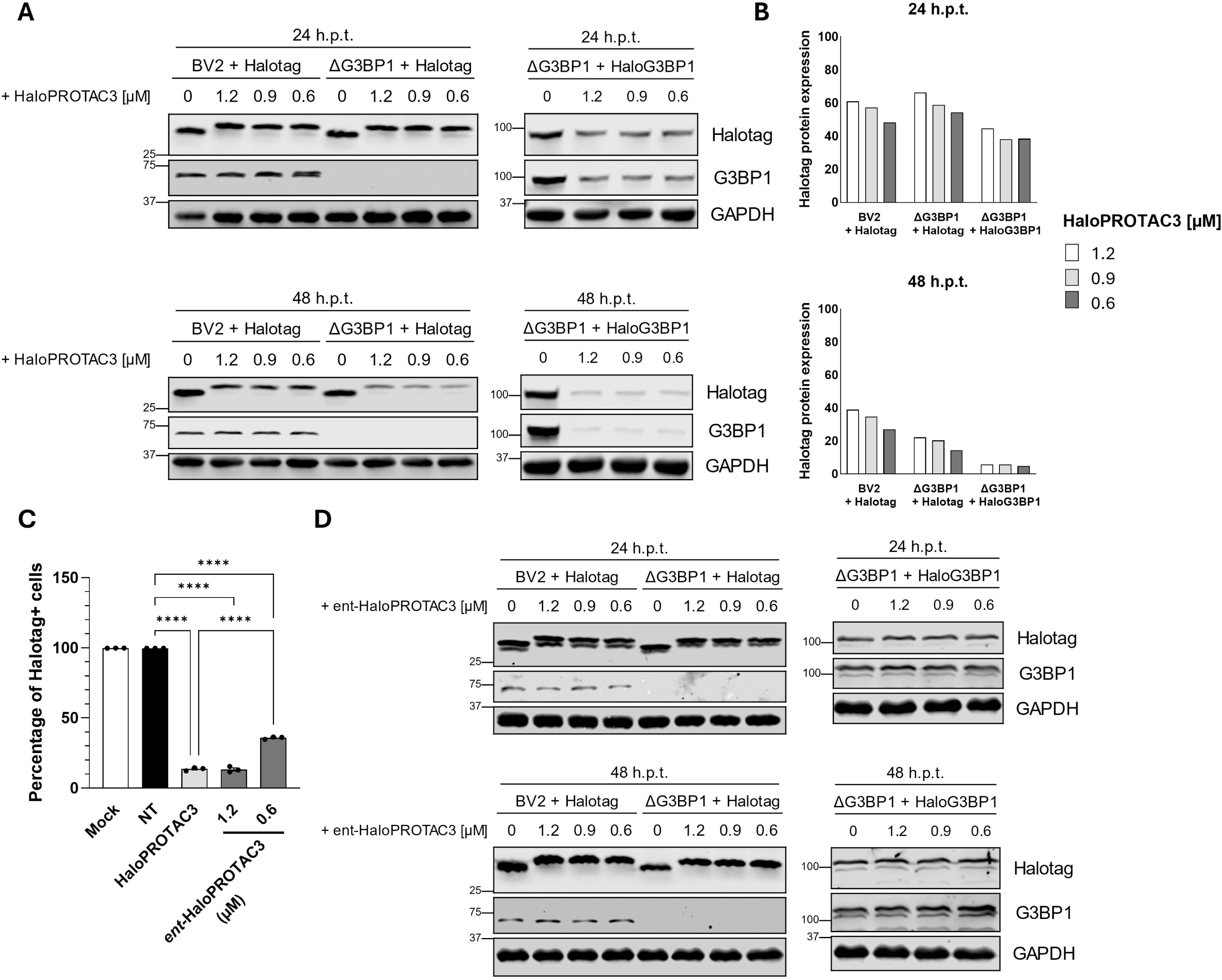
HaloPROTA3 degrades Halotag and HaloG3BP1 fusion proteins. (A-D) Protein samples from BV2 + Halotag, ΔG3BP1 + Halotag or ΔG3BP1 + HaloG3BP1 cells treated for either 24 h or 48 h with HaloPROTAC3 or *ent*-HaloPROTAC3 were subjected to western blot analysis using antibodies against Halotag, G3BP1 or GAPDH. (A) Representative western blots and (B) quantification of the Halotag protein expression in cells treated with HaloPROTAC3 normalised to GAPDH. (C) BV2 ΔG3BP1 + HaloG3BP1 cells treated for 48 h with either 1.2 µM of HaloPROTAC3 or the indicated concentrations of *ent*-HaloPROTAC3 were subjected to flow cytometry analysis as stated in Fig. 3C-E. (D) Representative western blots showing Halotag protein expression in cells treated with *ent*-HaloPROTAC3.

To confirm that the observed reduction in HaloG3BP1 protein levels was dependent on the interaction of HaloPROTAC3 with the VHL E3 ubiquitin ligase, an enantiomeric form of HaloPROTAC3 (*ent*-HaloPROTAC3, Figure 4C-D) was used in follow-up experiments. *Ent*-HaloPROTAC3 has a similar structure to HaloPROTAC3 but contains _D_-hydroxyproline and _D_-valine residue modifications (39) and is unable to bind VHL (Figure 3A). *Ent*-HaloPROTAC3 was able to engage all Halotag proteins effectively, resulting in a dose dependent reduction in Janelia 647 cells as assessed by FACS (Figure 4C). Notably however, the levels of the target protein remained unchanged as shown by western blot (Figure 4D). Binding was also evidenced as a shift in Halotag protein migration following SDS-PAGE (Figure 4D).

### HaloG3BP1 degradation with HaloPROTAC3 shows antiviral activity against MNV1

Having shown that HaloPROTAC3 efficiently degrades Halotag and HaloG3BP1, we then examined the antiviral effect of G3BP1 targeted degradation on norovirus infection. Given the time required for HaloPROTAC3 to degrade HaloG3BP1, the experimental strategy shown in Figure 5A was employed. Cells were treated for 48 h with 1.2 µM of the degrader prior to infection with MNV1. After a 1 h incubation to allow virus binding, HaloPROTAC3 was re-added and the infection allowed to continue for the indicated times, before samples were collected for analysis. The percentage of cells with detectable levels of the double-stranded RNA replication intermediate (dsRNA), a marker of active virus replication, was quantified by flow cytometry (Figure 5B-D). Alongside, Janeila 647 was used to determine the percentage of Halotag+ cells (Figure 5B-C). We observed a dose-dependent reduction in the percentage of HaloG3BP1 positive cells and a concomitant reduction in the number of cells with detectable levels of dsRNA, following treatment with HaloPROTAC3 (Figure 5B-D). We note however that a significant reduction in the number of dsRNA positive cells was only observed at concentrations > 0.6 µM. This data indicates that HaloG3BP1 targeted degradation using HaloPROTAC3 has an antiviral effect against MNV1 infection in the cellular model developed in this study.

**Figure 5.**
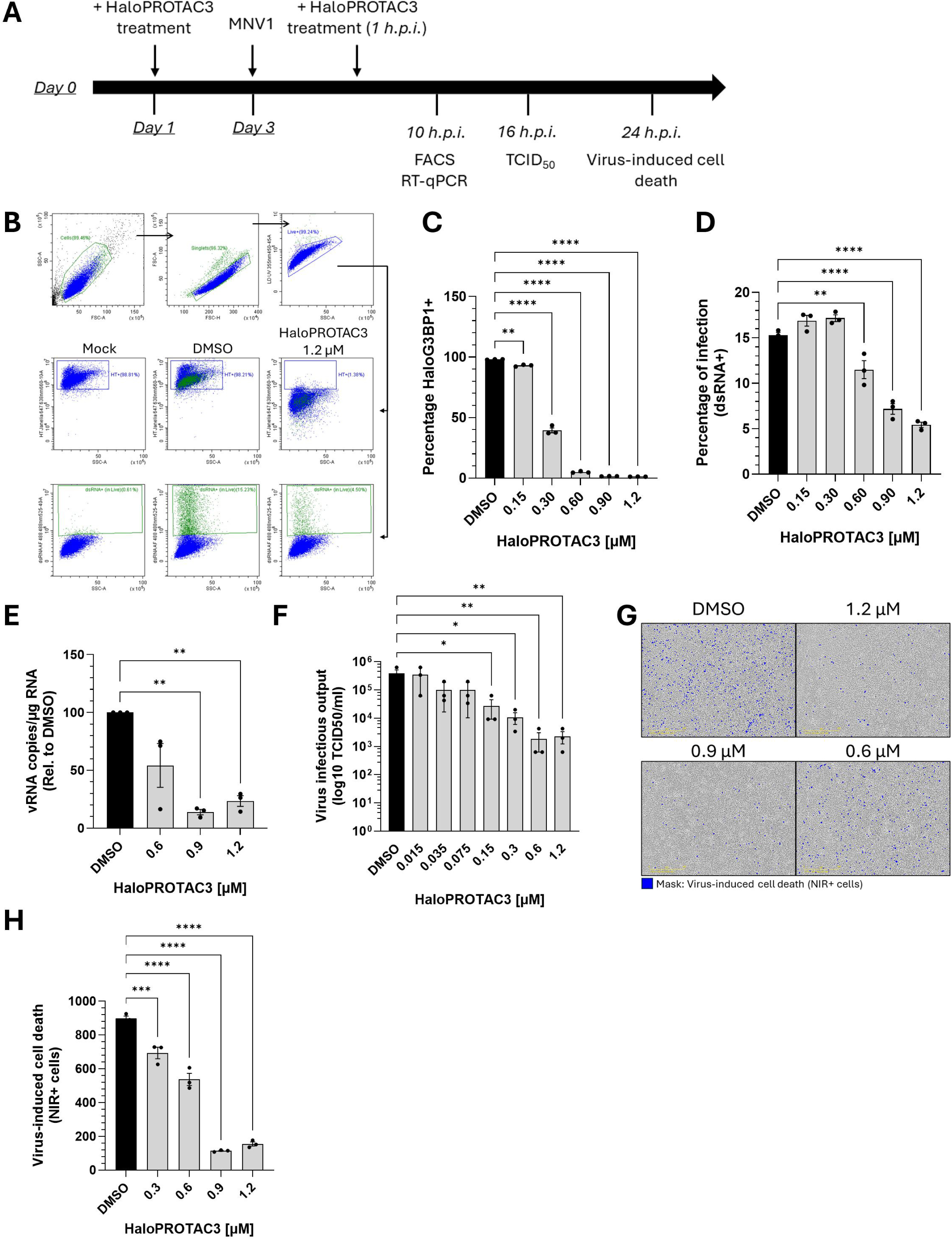
Degradation of G3BP1 using HaloPROTAC3 shows antiviral activity against norovirus. BV2 ΔG3BP1 + HaloG3BP1 cells were treated for 48 h with the shown concentrations of HaloPROTA3 and infected with MNV1. After 1 h of virus adsorption, virus inoculum was removed and HaloPROTAC3 was re-added to the cell culture medium for the remaining infection time. (A) Scheme of the experimental setup of the antiviral assays. (B-D) Cells infected with MOI 1 TCID_50_/cell for 16 h were incubated with fluorescent ligand Halotag Janelia 647 for 2 h, harvested and stained with anti-dsRNA antibody to assess the percentage of infection. Cells were then subjected to flow cytometry analysis. (B) Gating strategy of Halotag + cells and dsRNA + cells in the live cell population. (C) Quantification of the percentage of Halotag + and (D) dsRNA + (infected) cells across HaloPROTAC3 concentrations. (E) Total RNA was isolated from cells infected with MOI 1 TCID_50_/cell for 10 h and used to quantify the levels of viral RNA (expressed as vRNA copies/µg RNA, normalised to the DMSO control), by RT-qPCR. (F) Extracellular infectious virus yield in supernatants from cells infected with MOI 1 TCID_50_/cell for 16 h and treated with the indicated concentrations of HaloPROTAC3 was quantified by TCID_50_ in BV2 cells. (G-H) Infection-induced cell death was detected using the NIR dye measured in the live cell imaging Incucyte system. (G) Representative pictures after 24 h.p.i., NIR + cells are shown in blue. (H) Quantification of the total number of NIR + cells for the shown HaloPROTAC3 concentrations.

Our previous data indicated that G3BP1 likely plays a key role in norovirus VPg-dependent translation, but other roles in the viral life cycle have also been suggested (27–29). To further confirm our findings, and to elucidate if HaloPROTAC3 treatment has an effect on viral RNA synthesis, we quantified the levels of intracellular viral RNA by RT-qPCR. In agreement with the flow cytometry data, a statistically significant ∼80% decrease in the viral RNA copies was observed in cells treated with the highest concentration of HaloPROTAC3, but this effect was not significant at lower concentrations of the degrader (Figure 5E).

To further validate these observations, and to confirm that these effects translate to an impact on infectious virus yield, we quantified infectious virus released by infected cells by TCID_50_ after 16 h.p.i. (Figure 5F). In comparison to the DMSO-treated control cells, HaloG3BP1 knockdown using HaloPROTAC3 led to a significant dose-dependent reduction in the infectious virus titres. ΔG3BP1 + HaloG3BP1 cells treated with 1.2 µM HaloPROTAC3 exhibited a ∼99% decrease in virus released into the supernatant, with a similar reduction when cells were treated with 0.6 µM of the degrader (Figure 5F). Lastly, we assessed the virus-induced cell death in the presence of HaloPROTAC3 using the Incucyte imaging system. Figure 5G shows representative images taken after 24 h.p.i. in cells treated with DMSO or HaloPROTAC3. In agreement with our previous observations, treatment with 1.2, 0.9 and 0.6 µM of HaloPROTAC3 led to a statistically significant reduction in the number of dead cells (NIR+), indicating a protection of ΔG3BP1 + HaloG3BP1 cells from norovirus-induced cell death (Figure 5H).

### HaloPROTAC3 antiviral activity is mediated by binding and degradation of G3BP1

Given that the infectious virus yields from BV2 + Halotag and ΔG3BP1 + Halotag cells, remained unchanged after HaloPROTAC3 treatment (Figure S1), we concluded that HaloPROTAC3 has no indirect effects on norovirus replication and that the antiviral effect in ΔG3BP1 + HaloG3BP1 cells is likely driven by the degrader effect. However, to further confirm that the antiviral mechanism of action involved the proteasome ubiquitin system, we sought to examine if the effect required the engagement of the VHL-RING finger type E3 ubiquitin ligase. Binding of HaloPROTAC3 to Halotag fusion protein via the chloroalkane moiety is irreversible, which, in addition to inducing degradation, has the potential to affect the wider functions of G3BP1. We have shown in Figure 4 that, while *ent*-HaloPROTAC3 engages Halotag and the HaloG3BP1 fusion protein, it does not induce target degradation (Figure 4D). In line with this, we found that the infectious virus yield was unaffected (Figure 6A) by *ent*-HaloPROTAC3, as was virus-induced cell death *ent*-HaloPROTAC3 (Figure 6B). To explore the contribution of VHL engagement further, ΔG3BP1 + HaloG3BP1 cells were treated with HaloPROTAC3 in the presence and absence of VH032, a VHL inhibitor that binds to the same binding site as HaloPROTAC3 and should compete for its interaction with VHL. We first ensured that the concentrations of VH032 were sub-toxic, especially when used in combination with HaloPROTAC3 (Figure 6C). As expected, we observed that treatment with HaloPROTAC3 in the presence of increasing concentrations of VH032 led to a dose-dependent recovery in HaloG3BP1 protein levels, particularly after 48 h.p.t. (Figure 6D). We noted however, that the degradation of HaloG3BP1 protein was not fully prevented (Figure 6D), likely due to VH032 having lower affinity for VHL in comparison to VH285, the VHL-binding warhead on which HaloPROTAC3 is based (39). Furthermore, we also observed that VH032 was able to partially restore virus-induced cell death (Figure 6E) and infectious viral yield from cells treated with HaloPROTAC3 in the presence VH032 (Figure 6F). VH032 treatment did not show any effect in virus replication when used as a single treatment (Figure S2).

**Figure 6.**
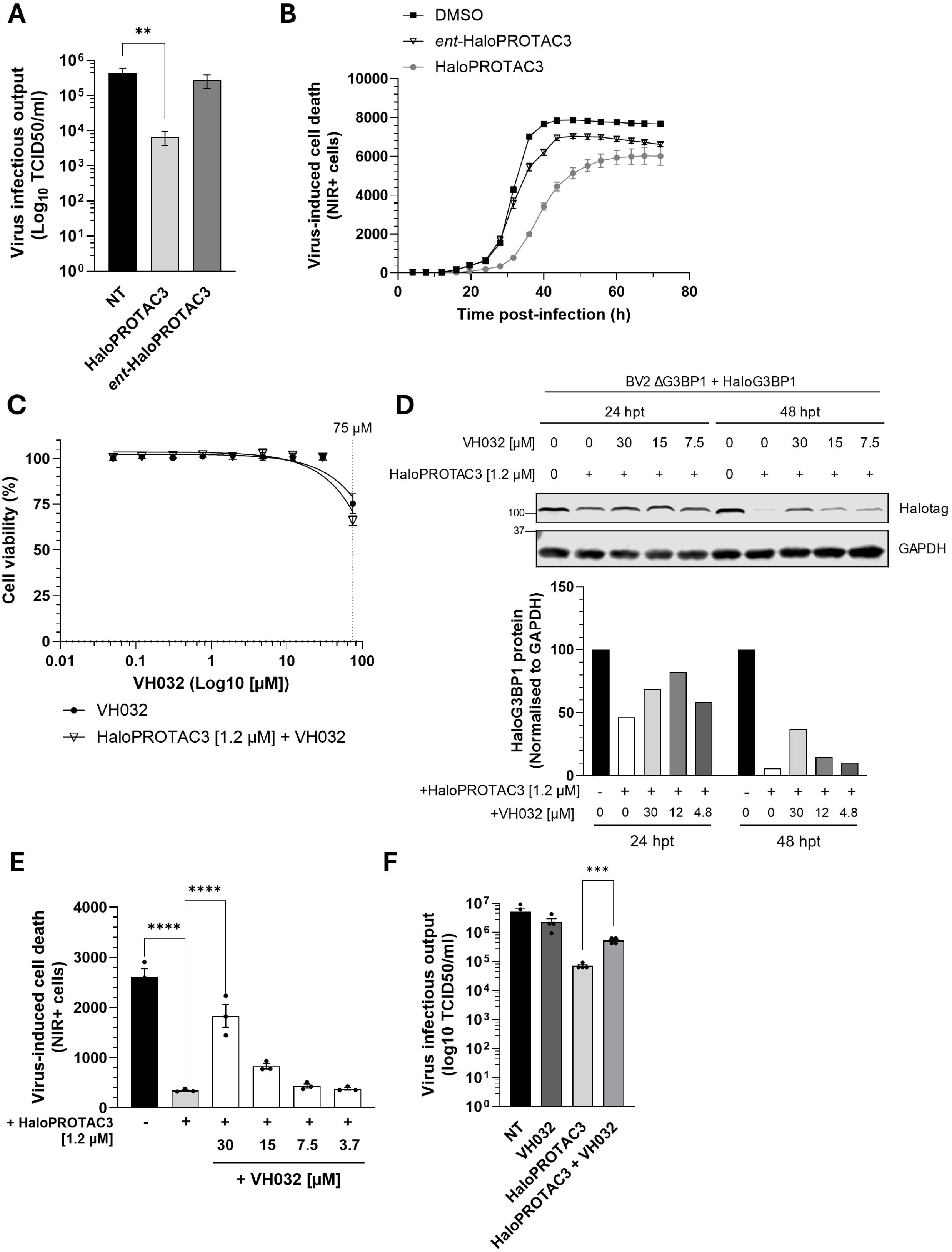
Degradation of G3BP1 is required for HaloPROTAC3 antiviral activity. (A-B) BV2 ΔG3BP1 + HaloG3BP1 cells were treated for 48 h with 1.2 µM of HaloPROTAC3 or *ent*-HaloPROTAC3 as indicated and infected with MNV1. After 1 h of virus adsorption, virus inoculum was removed and PROTACs re-added to the cell culture medium. (A) Extracellular infectious virus yield in supernatants from cells infected with MOI 1 TCID_50_/cell for 16 h and treated with either HaloPROTAC3 or *ent*-HALOPROTAC3 was quantified by TCID_50_ in BV2 cells. (B) Infection-induced cell death was detected using the NIR dye measured in the live cell imaging Incucyte system. Shown is the quantification of the total number of NIR + cells for cells treated with either HaloPROTAC3 or *ent*-HaloPROTACs, measured every 3 h and followed for a total of 72 h.p.i. (C-F) BV2 ΔG3BP1 + HaloG3BP1 cells were concomitantly treated for 48 h with 1.2 µM of HaloPROTAC3 and VH032 in the indicated concentrations. (C) Metabolic cell death was measured using Cell titer blue. Cell death was calculated as relative to the DMSO control. 75 µM of VH032 is indicated by a dotted line. (D) Protein samples from cells treated for either 24 or 48 h with the indicated concentrations of HaloPROTAC3 and VH032 were subjected to western blot analysis using antibodies directed Halotag or GAPDH. Bar plot shows the quantification of the Halotag protein bands for all the experimental conditions. (E) Cells were infected with MOI 1 TCID_50_/cell of MNV1. After 1h of virus adsorption, virus inoculum was removed and drugs re-added to the cell culture medium. Infection-induced cell death was detected using the NIR dye measured in the live cell imaging Incucyte system as stated in (B). Shown is the quantification of the total number of NIR + cells measured after 24 h.p.i. (F) Extracellular infectious virus yield quantified by TCID_50_ in BV2 cells from supernatants of cells infected with MOI 1 TCID_50_/cell for 16h and treated with 1.2 µM of HaloPROTAC3 and 30 µM of VH032.

Overall, we have demonstrated that HaloPROTAC3-mediated degradation of HaloG3BP1 in the cellular model developed in this study is dependent on the proteasome-ubiquitin system via its interaction with Cullin2-VHL-RING finger type E3 ubiquitin ligase. Furthermore, this interaction is required for the antiviral activity exhibited by HaloPROTAC3.

## Discussion

Despite its clinical significance, particularly in high-risk group individuals, HNoV remains a pathogen for which no specific antiviral therapeutics are currently available. Host-targeted antiviral therapeutics is emerging as a promising approach to overcome common obstacles in RNA virus drug discovery (40, 41). In this study, we explored the antiviral potential of targeted protein degradation of the host factor G3BP1, which is required for both human and murine norovirus infection. We provide a proof of concept that degradation of G3BP1 using a PROTAC impairs norovirus replication and therefore constitutes a potential novel antiviral strategy.

Since HNoV cultivation *in vitro* remains a challenge, we used the well-characterised MNV1 model to establish a genetic system enabling the conditional degradation of G3BP1. We generated stable cells in which G3BP1 was genetically ablated and reconstituted with G3BP1 fused to Halotag, a bacterial protein that covalently binds chloroalkane-based synthetic ligands (42). Reconstitution of BV2 ΔG3BP1cells with a Halotag-G3BP1 fusion protein restored virus replication, validating both the proviral function of G3BP1 and the suitability of this fusion protein for functional complementation. The cellular platform developed here enabled us to induce the specific proteolysis of G3BP1 using the heterobifunctional HaloPROTAC3 that incorporates a VHL ligand (VH287) and a chloroalkane warhead for Halotag engagement (39). This exploratory approach allowed us to circumvent the lack of specific G3BP1-directed PROTACs at the time when this study was undertaken.

Treatment with 1.2 µM HaloPROTAC3 for 48 h resulted in maximal target degradation of HaloG3BP1 (95.3%), compared to 85.8% and 72.9% in ΔG3BP1 + Halotag and WT cells, respectively. These levels are in line with previous reports using HaloPROTACs for the degradation of endosomal and cytosolic proteins (38, 39). Our estimated DC₅ ₀ (∼600 nM) is noticeably higher than the 19 nM reported previously in other studies (39). Variations in degradation potency in between studies likely reflect differences in the method and mammalian promoters used for over-expression, intrinsic cell-specific variables or the method used as a readout. Notably, antiviral activity closely paralleled protein degradation: maximal antiviral effect was observed at 1.2 µM, with significant effects retained at 150 nM. Degradation of G3BP1 reduced intracellular MNV1 RNA levels and infectious virus yield and also protected cells from virus-induced cytopathic effect.

Host-targeted antiviral PROTACs remain relatively underexplored, although a number of notable examples have been reported. Indomethacin-derived PROTACs, for example, have been developed against coronaviruses (36). The mechanism of action was suggested to be based on the degradation of human prostaglandin E synthase type 2 (PGES-2), a protein that plays an important role in coronavirus replication by binding to NSP7 virus protein during replication (36). However, this mode of action and the degradation efficiency of these compounds needs further exploration. More recently, Newton, et al., designed PROTACs targeting human protein cyclophilin A, a host cofactor for both human immunodeficiency virus and hepatitis C virus (HCV) (37). On the other hand, most of the antiviral PROTACs developed to date correspond to the class of direct-acting antivirals, and have been designed with the aim of targeting viral proteins (43–48). PROTACs targeting HCV serine protease (NS3/4A) using the antiviral small-molecule inhibitor telaprevir as a ligand, were shown to significantly induce proteasomal degradation of the viral target (43). The clinically approved SARS-CoV-2 Mpro inhibitor nirmatrelvir was similarly used to develop potent antiviral degraders, utilising VHL or IAP ubiquitin ligase recruiters (48). The neuraminidase inhibitor oseltamivir was also used for the design of PROTACs against influenza A virus (IAV), showing good anti-H1N1 activity and efficient IAV neuraminidase degradation (46). IAV-directed PROTACs have been developed to also target the hemagglutinin (HA) virus protein. Li et al., showed that oleanolic acid derivatives conjugated to VHL ligands were able to bind and deplete HA with a DC_50_ value of 1.44 µM (47). Collectively, these studies have demonstrated that targeted protein degradation overcomes challenges of traditional small molecule inhibitors such as drug resistance emergence and virus diversity. Indeed, PROTAC-mediated antiviral activity against oseltamivir-resistant IAV and nirmatrelvir-resistant SARS-CoV-2 strains has been proven (46, 48).

We confirmed that HaloPROTAC3-mediated G3BP1 degradation and antiviral activity depends on VHL recruitment. The enantiomeric form of the degrader, unable to bind VHL, did not result in target degradation or norovirus inhibition. Similarly, co-treatment with the VHL inhibitor VH032 blocked G3BP1 depletion and reversed the antiviral phenotype. These findings are consistent with previous work using VHL inhibitors or proteasome blockade to demonstrate the dependency of HaloPROTAC activity on E3 ligase recruitment and proteasomal degradation (38, 39).

Due to its central role in stress granules assembly (49, 50), G3BP1 is commonly hijacked by both DNA and RNA viruses to support their replication (51). In the case of noroviruses, at least one function of G3BP1 is to facilitate the recruitment of ribosomal subunits to VPg-linked viral RNA, thus promoting viral protein synthesis (27). A similar strategy was previously reported for several viruses from the alphavirus genus (52). G3BP1 was additionally found to associate with the norovirus replication complex (28). Similarly, infections with Zika virus (ZIKV) and alphaviruses like chikungunya virus (CHIKV) result in G3BP1 clustering within the virus replication complex, driven by interactions between this host factor and viral proteins (52–54). This can enhance virus replication by suppressing the host antiviral response, more specifically, by inhibiting stress granules assembly (52–54). Given that viruses from diverse taxonomic families rely on G3BP1 function, this emerges as a promising target for broad-spectrum antiviral therapeutics which should be explored further. Recently, a newly synthesized compound was used for the development of PROTACs targeting G3BP1/2 (55). These degraders used pomalidomide analogues for CRBN E3 ligase recruitment coupled with a PEG3 linker to the G3BP1/2-targeting moiety. The authors achieved a maximum degradation of 95% of G3BP1 and G3BP2 at concentrations of 5 µM and 10 µM respectively (55), and the degradation was abrogated by proteasome inhibition with MG132 (55). Further research is necessary to elucidate whether this PROTAC possess antiviral activity against norovirus and potentially other RNA viruses. Likewise, small molecules targeting the nuclear transport factor 2-like domain of G3BP1 that were recently identified and shown to have antiviral activity against CHIKV (56), could be potentially used for the rational design of G3BP1-targeting degraders. In this way, therapies targeting G3BP1 could be combined with direct-acting antivirals to enhance their efficacy and reduce the risk of resistance emergence. However, since G3BP1 has an essential physiological role, its rapid degradation may involve unknown risks. Therefore, future studies on the use of PROTACs directed towards G3BP1 or other host factors, should focus on assessing feasibility and safety.

Although a lot of progress has been made into the development of PROTACs, with various in Phase 2 and 3 clinical studies, particularly for cancer treatment, there are still a lot of challenges in the field (57). *In vivo* stability and efficient delivery to the target cells is a common concern that has been addressed using nanoparticle-based delivery systems (58). As mentioned above, the potential use of PROTACS is hindered by the possibility of off-target effects, hence, strategies to refine specificity need to be put in place to minimise cytotoxicity due to unintended degradation of other proteins (59).

In summary, we demonstrate that targeted protein degradation of G3BP1 via HaloPROTAC3 provides a tractable approach to impair norovirus replication. This platform validates the use of conditional degrader systems to probe host-virus interactions and highlights G3BP1 as a promising target for future host-directed antiviral interventions.

## Materials and Methods

### Compounds

HaloPROTAC3 (VH285-PEG4-C4-Cl, HY-111997) and VH032 (HY-120217) were purchased from MedChemExpress. *ent*-HaloPROTAC3 was obtained from Promega. All compounds were reconstituted in DMSO and used at the indicated concentrations. Final DMSO concentration did not exceed 0.05%, unless stated otherwise. Resazurin stock solutions were prepared by dissolving in phosphate-buffered saline (PBS) at a concentration of 5 mg/mL and filtering through a 0.45 µm filter.

### Cells

All the cell lines used in this study were cultured in complete Dulbecco’s Modified Eagle’s Medium (DMEM) 4,5 g/L D-glucose, sodium bicarbonate and sodium pyruvate. Medium was supplemented with 10% heat-inactivated Foetal Calf Serum (HyClone), 2 mM L-glutamine, MEM non-essential amino acids (Sigma-Aldrich) and antibiotics (10U/ml of penicillin and 100 μg/mL of streptomycin). The murine microglial BV2 cell line (60) was a kind gift from Dr. Jennifer Pocock (University College London), HEK293T were kindly provided by Dr. Susanna M. Colaco (University of Cambridge). G3BP1 knockout (ΔG3BP1) BV2 cells clone 1B2, were previously described (27). BV2 + Halotag, BV2 ΔG3BP1 + Halotag and BV2 ΔG3BP1 + Halotag G3BP1 were all generated by lentivirus-mediated transduction (see below) and maintained in medium containing 400 µg/mL Hygromycin B (Invitrogen). Baby Hamster Kidney (BHK) cells expressing the T7 RNA polymerase (BSR-T7) were obtained from Karl-Klaus Conzelmann (Ludwid Maximillians University, Munich) and maintained in medium supplemented with 0.5 mg/mL G418 (Invivogen). Cells were all incubated at 37°C with 5% CO_2_ and screened monthly for mycoplasma and confirmed as negative.

### Plasmids

The full open reading frame of the human G3BP1 sequence fused at the N-terminal to the bacterial dehalogenase Halotag (pFN21AAA0660 Halotag CMV Flexi vector, Promega) was subcloned into a lentiviral 3^rd^ generation transfer plasmid using SgfI and PmeI restriction enzymes (Promega). An additional lentivirus plasmid encoding Halotag was generated by addition of a stop codon (TAA) at the 3’ of the Halotag sequence, by PCR amplification with KOD Hot Start DNA polymerase enzyme (Merk Sigma-Aldrich), using the original vector as template. Sequences of all the generated constructs were verified by Oxford Nanopore sequencing (Department of Biochemistry, University of Cambridge).

### Lentivirus production and transduction

Infectious transgenic lentiviruses were produced in HEK293T cells by co-transfection of the transfer plasmids (Halotag-G3BP1 or Halotag), together with packaging (pMDLg/pRRE, pRSV-Rev) and envelope (pMD2.G) plasmids, using Lipofectamine 2000 (Invitrogen) following manufacturer’s recommendations. Lentivirus-containing supernatants were harvested after 2 days post-transfection and used to infect BV2 WT or ΔG3BP1 cells. Subsequently, stable cells were selected with 400 μg/mL Hygromycin B (Invitrogen) for two weeks. Cells were stained with the fluorescent Janelia Fluor 646 Halotag ligand (Promega, GA1120) diluted in culture medium at a final concentration of 200 nM for 2 h, and high Halotag-expressing cells were enriched by sorting in a BD FACSAria III Cell Sorter (BD Biosciences).

### Virus production and titration

MNV1 was produced by reverse genetics in BSR-T7 cells as described previously (61). Briefly, cells were infected with Fowlpox virus (FPV)-T7 at an approximate multiplicity of infection (MOI) of 0.5 PFU/cell for 2 h and subsequently transfected with pT7:MNV-1_3′Rz cDNA infectious clone (62) using Lipofectamine 2000 (Invitrogen), following manufacturer’s instructions. Virus was harvested after 2 days post-infection (d.p.i.), freeze-thawed and titrated by TCID_50_. MNV1 high titre virus stocks were then produced by infecting BV2 WT cells at an MOI of 0.01 TCID_50_/cell. Cells were incubated for 24 h, until ∼50% CPE was reached. Virus-containing supernatant was freeze-thawed once, clarified at 1000 x g for 10 min at 4°C to remove cell debris and filtered through a 0.22 µm filter.

Virus titres were estimated by determination of the TCID_50_/mL in BV2 cells. Briefly, cells were plated in 96 well plates and infected with 10-fold serial dilutions of the virus samples taken at the specified time-points. After 2 d.p.i., TCID_50_ endpoints were determined by scoring CPE in the infected cells and viral titres calculated using the Reed-Muench method (63).

### Cytotoxicity assays

Cell toxicity induced by all the drugs used in this study was quantified using the resazurin reduction assay as previously described (64). In brief, cells were plated in 96 well plates 24 h prior treatment, subsequently, cells were treated with the indicated concentrations of either HaloPROTAC3, *ent*-HaloPROTAC3 or VH032 (as a single treatment or in combination with HaloPROTAC3), all diluted in culture medium. After 48 h the medium was removed and replaced by culture medium containing resazurin at a final concentration of 100 µg/mL. Plates were placed back in the incubator for 4 h and fluorescence at 560nm/590nm was measured on a SpectraMax i3 plate reader (Molecular Devices). Cell death is expressed as percentage in comparison to the non-treated or the DMSO control cells as stated in the text.

### Virus-induced cell death assays

To assess virus-induced cell death in an end-point assay, BV2 WT, BV2 + Halotag, BV2 ΔG3BP1 + Halotag or BV2 ΔG3BP1 + Halotag G3BP1 cells were seeded in 96-well plates 24 h prior to infection with 10-fold serial dilutions of MNV1. After 5 d.p.i., cells were fixed with formal saline (4% formaldehyde v/v) and stained with toluidine blue prior to scanning.

Live cell imaging was employed to monitor and quantify virus-induced cell death in real-time. Cells were seeded in 96-well plates one or two days prior to infection with MNV1 at the indicated MOI. For the antiviral assays, cells were plated and treated with HaloPROTAC3 two days prior to infection. After 1 h of virus adsorption at 37°C, the inoculum was removed and replaced by complete medium containing Incucyte Cytotox near infrared (NIR) Dye (Sartorius) at a final concentration of 0.6 µM, in the presence or absence of HaloPROTAC3. Plates were placed in the Incucyte live-cell imaging system and images were captured at the defined time intervals stated in the figure legends, with the 10X objective and in the phase contrast and NIR channels. Each condition was evaluated in triplicate and at least five pictures were taken per well. The NIR Object count, indicating cell death events, was quantified using Incucyte Basic Analysis Software version 2023A Rev1 and expressed as NIR positive (NIR + cells) per condition.

### Quantification of MNV1 RNA copies by RT-qPCR

RNA was extracted from infected cells using GenElute Mammalian Total RNA Miniprep Kit (Sigma-Aldrich) according to the manufacturer’s protocol. DNA removal from RNA samples was done with DNase I (Invitrogen) and cDNA was synthesized using M-MLV Reverse Transcriptase (Promega) and random hexamers (Merck-SigmaAldrich). SYBR Green based qPCR was carried out using a master mix containing 2.5mM MgCl_2_, 400 μM dNTPs, 1/10,000 SYBR Green (Molecular Probes), 1 M Betaine (Sigma), 0.05 U/μl of Gold Star polymerase (Eurogentec), 1/5 10X Reaction buffer (750 mM Tris-HCl pH 8.8, 200 mM [NH4]_2_SO_4_, 0.1% [v/v] Tween 20, without MgCl_2_), and ROX Passive Reference buffer (Eurogentec). qPCR reactions were run on a ViiA 7 Real-Time PCR System (ThermoFisher Scientific). Absolute MNV1 RNA copies/mL in each sample were calculated using the standard curve method. The standard was prepared with 10-fold serial dilutions of pT7:MNV-1_3′Rz cDNA infectious clone (62).

### Western blot

Cells treated with the compounds used in this study and either infected with MNV1 or mock-infected, were washed with PBS and lysed in RIPA buffer (150mM NaCl, 1% NP-40, 0.5% Na deoxycholate, 0.1% SDS, 25mM Tris-HCl pH 7.4) supplemented with Halt Protease and Phosphatase Inhibitor Cocktail (Thermo Scientific). Cells were lysed at 4°C, on rotation, for 20 min. Protein lysates were clarified by centrifugation at 10,000 x g for 10 min at 4°C. Protein concentrations were quantified using Pierce BCA assay (Thermo Scientific) according to the manufacturer’s recommendations. SDS sample buffer (0.0625M Tris-Cl pH 6.8, 10% glycerol, 5% 2-mercaptoethanol, 2% SDS, 0.002% bromophenol blue) was used to prepare the samples, which were subsequently heated at 95°C for 5 min, prior to separation by SDS PAGE in acrylamide Tris-Glycine gels. Proteins were transferred onto 0.45 μm nitrocellulose membranes and blocked with 5% milk for 1 h at room temperature, prior to incubation with primary antibodies at 4°C, overnight. Primary antibodies used in this study were as follows: mouse anti-GAPDH mAb (6C5) (Invitrogen, AM4300), rabbit anti-Halotag pAb (Promega, G9281), rabbit anti-G3BP1 pAb (ARP37713_T100, Aviva Systems Biology), rabbit anti MNV1 NS7 pAb (non-commercial). Membranes were subsequently incubated for 1 h at room temperature with the secondary antibodies IRDye 680RD Donkey anti-Mouse IgG and IRDye 800CW Goat anti-Rabbit IgG (LICORBio). Western blots were scanned using Odyssey CLx imager (LICORBio) and the results were analysed and quantified using Image Studio Lite software version 5.2.5 (LICORBio).

### Flow cytometry

The expression of Halotag (or Halotag G3BP1) was examined by FACS analysis. BV2 WT, BV2 + Halotag, BV2 ΔG3BP1 + Halotag or BV2 ΔG3BP1 + Halotag G3BP1 were seeded in 24 well plates and treated with the specified concentrations of HaloPROTAC3 or *ent*-HaloPROTAC3. After 48 h post-treatment (h.p.t.), medium was removed and cells were incubated for 2 h with the fluorescent Janelia Fluor 646 Halotag ligand (Promega, GA1120), diluted in culture medium at a final concentration of 200 nM. After the incubation time, the cells were detached from the plates by using 0.25% Trypsin-EDTA solution (Sigma-Aldrich) and washed with ice-cold FACS Buffer (PBS without magnesium and calcium salts [PBS Mg_2_^-^/Ca_2_^-^], supplemented with 0.5% FCS, 2 mM EDTA, and 0.1% NaN_3_). Subsequently, cells were stained using LIVE/DEAD™ Fixable Blue Dead Cell Stain Kit, for UV excitation (Invitrogen) for 20 min at room temperature and fixed using Paraformaldehyde, 4% (w/v) aqueous solution, methanol free (Thermo Scientific) for 10 min at room temperature. In samples were the percentage of infection was also examined, the cells were washed and permeabilised for 10 minutes on ice with FACS buffer supplemented with 0.3% Triton X-100, and stained for 2 h with Mouse anti-dsRNA mAb (J2) (Cell Signaling, 76651). Cells were washed with FACS buffer and stained with Goat anti-Mouse IgG (H+L) Cross-Adsorbed Secondary Antibody, Alexa Fluor 488 (Invitrogen). Subsequently, the cells were subjected to analysis in a CytoFlex LX Flow Cytometer (Beckman Coulter) using the Blue 488nm (80mW), Red 638nm (50mW) and UV 355nm (20mW) lasers; and 525/40, 660/10, 450/45 filters; for detection in the green, red and blue spectra respectively. Data was analysed and visualised in the CytExpert software version 2.5 (Beckman Coulter).

### Statistical analysis

All the data in this work is presented as mean ± SEM from at least 3 independent biological experiments. Data visualisation and analysis was performed in GraphPad Prism version 10.4.1. Either *T*-student test or one-way ANOVA with Bonferroni multiple comparisons tests was applied to determine statistical significance, unless indicated otherwise. In all cases, ^∗^, ^∗∗^, ^∗∗∗^, and ^∗∗∗∗^ are used to denote p ≤ 0.05, p ≤ 0.01, p ≤ 0.001 and p ≤ 0.0001 respectively. Snapgene 4.2 was used for sequence alignments, cloning strategies and primer design.

## Acknowledgements

This research was funded by the Wellcome Trust grant 207498/Z/17/Z. We thank Dr. V. Lulla (University of Cambridge) for providing access to the Incucyte live-cell imaging system. The authors acknowledge the flow cytometry facility from the School of the Biological Sciences of the University of Cambridge for their assistance in this work.

## Author contributions

Liliana Echavarria-Consuegra: Conceptualization, Formal analysis, Investigation, Methodology, Visualisation, Writing – original draft, Writing – review & editing. Ian Goodfellow: Conceptualization, Funding acquisition, Project administration, Supervision, Writing – review & editing.

## Declaration of competing interest

None to declare.

## Supplementary data

**Figure S1.**
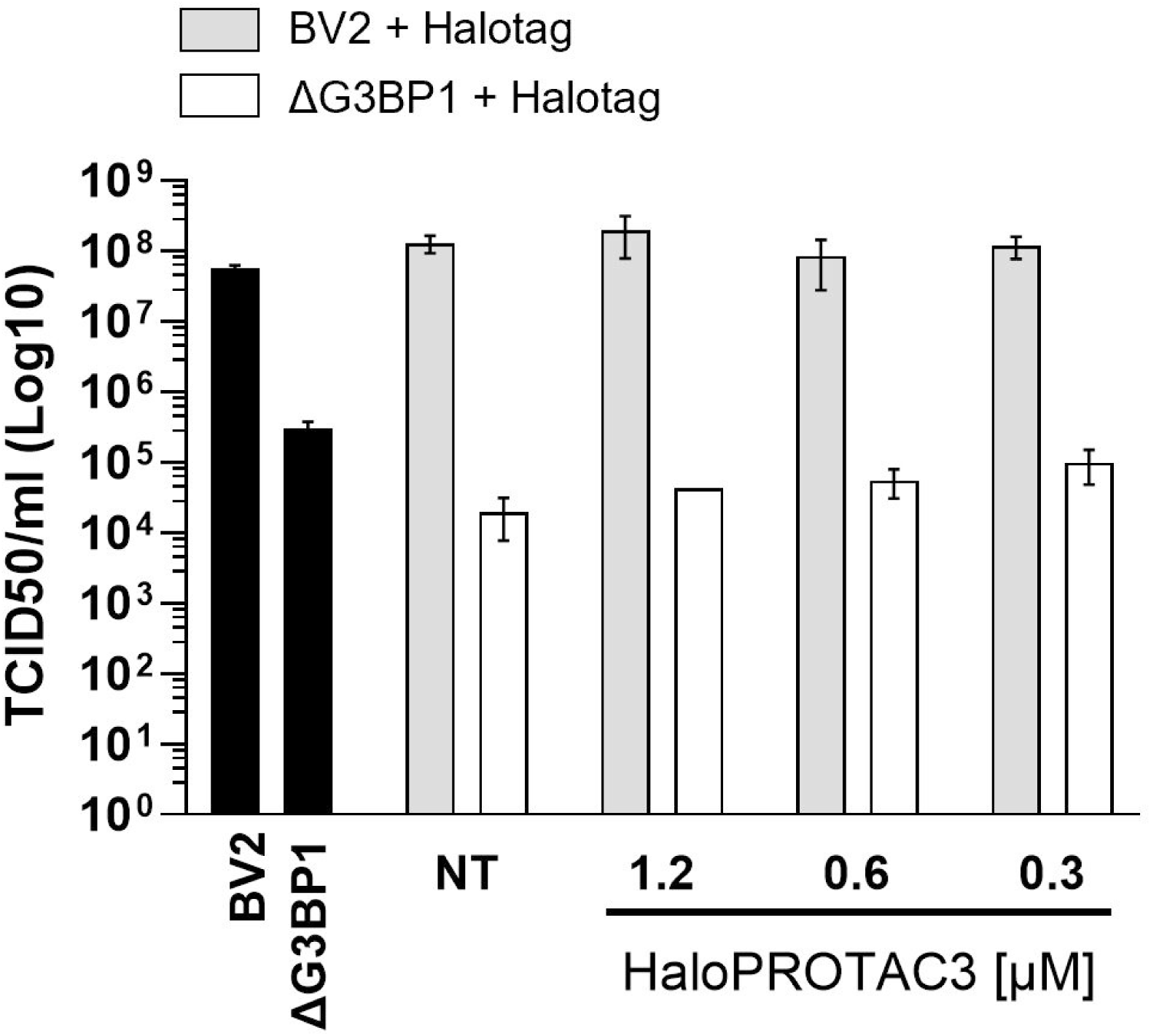
Extracellular infectious virus yield quantified by TCID_50_ in BV2 cells from supernatants of BV2, ΔG3BP1, BV2 + Halotag or ΔG3BP1 + Halotag cells infected with MOI 1 TCID_50_/cell and treated with the indicated concentrations of HaloPROTAC3.

**Figure S2.**
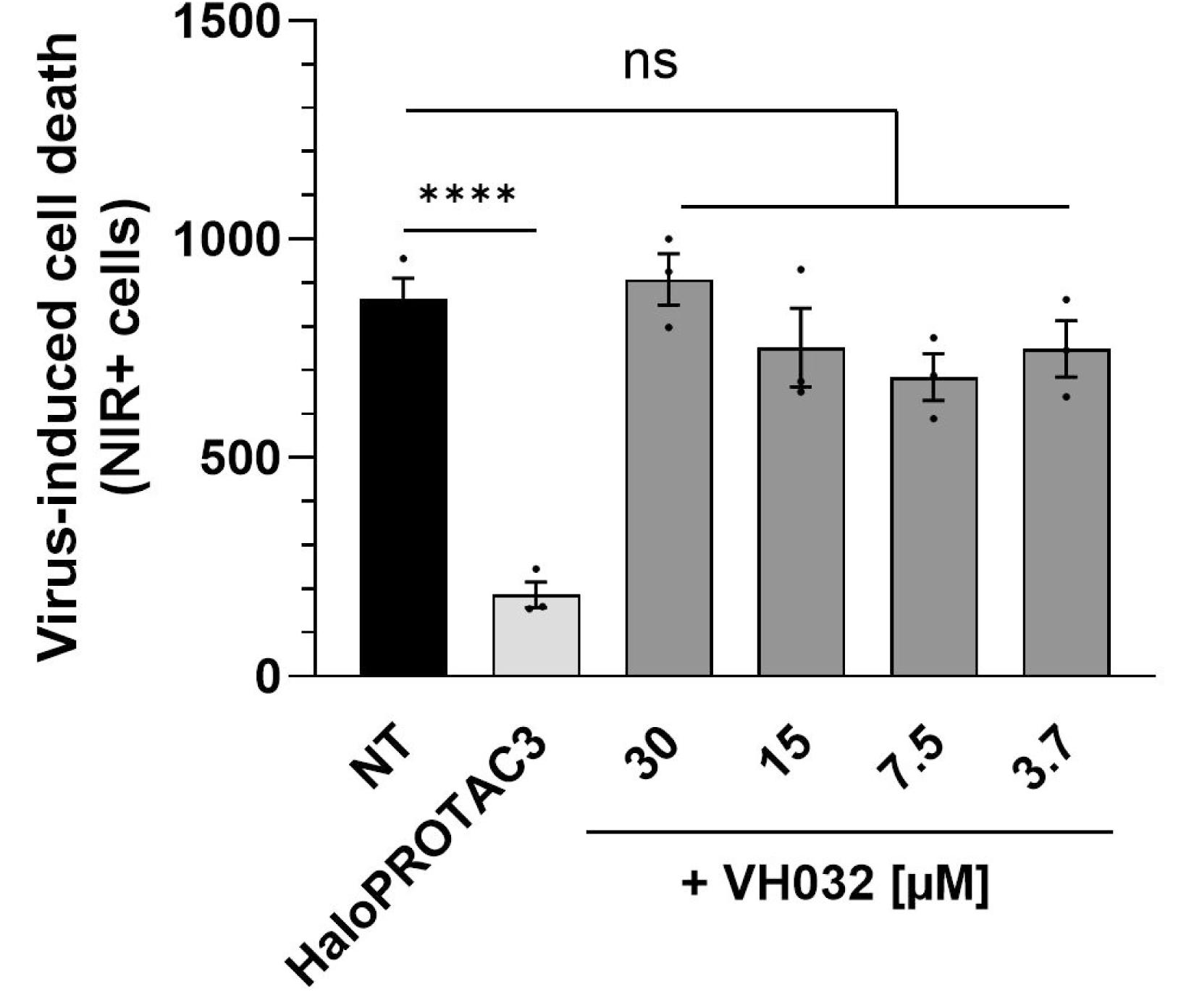
BV2 ΔG3BP1 + HaloG3BP1 were treated with either HaloPROTAC3 or VH032 at the indicated concentrations and infected with MNV1 at MOI 1 TCID_50_/cell. Infection-induced cell death was detected using the NIR dye measured in the live cell imaging Incucyte system. The quantification of the total number of NIR + cells after 24 h.p.i. is shown.

